# Scrutinizing the Feasibility of Macroscopic Quantum Coherence in the Brain: A Field-Theoretical Model of Cortical Dynamics

**DOI:** 10.1101/2023.03.03.530961

**Authors:** Joachim Keppler

**Affiliations:** Department of Consciousness Research, DIWISS, 91154 Roth, Germany

**Keywords:** brain dynamics, quantum field theory, zero-point field, collective behavior, phase transitions, criticality, coherence domains

## Abstract

The neural activity patterns associated with advanced cognitive processes are characterized by a high degree of collective organization, which raises the question of whether macroscopic quantum phenomena play a significant role in cortical dynamics. In order to pursue this question and scrutinize the feasibility of macroscopic quantum coherence in the brain, a model is developed regarding the functioning of microcolumns, which are the basic functional units of the cortex. This model assumes that the operating principle of a microcolumn relies on the interaction of a pool of neurotransmitter (glutamate) molecules with the vacuum fluctuations of the electromagnetic field, termed zero-point field (ZPF). Quantitative calculations reveal that the coupling strength of the glutamate pool to the resonant ZPF modes lies in the critical regime in which the criterion for the initiation of a phase transition is fulfilled, driving the ensemble of initially independent molecules toward a coherent state and resulting in the formation of a coherence domain that extends across the full width of a microcolumn. The formation of a coherence domain turns out to be an energetically favored state shielded by a considerable energy gap that protects the collective state against thermal perturbations and entails decoherence being greatly slowed down. These findings suggest that under the special conditions encountered in cortical microcolumns the emergence of macroscopic quantum phenomena is feasible. This conclusion is further corroborated by the insight that the presence of a coherence domain gives rise to downstream effects which may be crucial for the cortical communication and the formation of large-scale activity patterns. Taken together, the presented model sheds new light on the fundamental mechanism underlying cortical dynamics and suggests that long-range synchronization in the brain results from a bottom-up orchestration process involving the ZPF.

## I. INTRODUCTION

One of the major scientific challenges in the field of biological physics consists in unraveling the fundamental mechanisms that govern the brain dynamics of highly developed organisms. This applies particularly to the dynamics associated with advanced cognitive functions, especially those related to conscious processes, which are characterized by synchronized neural activity patterns extending over large cortical areas. These activity patterns originate from a vast number of neurons exhibiting collective behavior [1–4]. The body of empirical evidence that has been accumulated in recent years suggests that pattern formation is based on phase transitions [2], supporting the hypothesis that criticality underlies the organization of the brain [5–7]. However, the empirical data on their own do not provide insight into the details of the mechanism that leads to criticality [5, 6].

In view of these findings, it has been acknowledged that the tools used in theoretical physics are essential for a deeper understanding of the dynamical characteristics of a many-body system such as the brain [8]. Among these tools, approaches from quantum field theory have proven particularly useful in describing collective modes of a system and interpreting coherent activity patterns as macroscopic features of quantum origin, thus shedding light on the basic principles that may account for the formation of spatially extended domains of synchronized activity [9–11]. Furthermore, it has been pointed out that a consistent interpretation of the neural correlates of consciousness and a look behind the scenes of conscious processes is achievable by resorting to quantum electrodynamics (QED) [12–15]. According to this QED-based approach, the brain is postulated to function as a resonant oscillator that couples to the ever-present vacuum (zero-point) fluctuations of the electromagnetic field, which in the following will be referred to as zero-point field (ZPF). In this model, the ZPF plays a central role in the orchestration of brain activity and the formation of coherent activity patterns, opening up new perspectives for the development of a self-consistent theory of consciousness [15].

Even though these approaches are suitable for making qualitative statements about the physical principles driving brain activity, their acceptance depends on the provision of quantitative calculations using realistic, i.e., empirically backed, parameter values. This step is crucial for demonstrating the plausibility of the postulated mechanisms, particularly in view of the fact that the significance of macroscopic quantum phenomena in explaining neural activity patterns contradicts the common belief that classical physics should be sufficient to account for all aspects of brain activity [16]. For this reason, there is an urgent need to advance the existing models.

The aim of this paper is to scrutinize the feasibility of macroscopic quantum coherence in the brain. For this purpose, a model of the functioning of microcolumns, which constitute the basic functional units of the cor-tex and support high-level cognitive processes, is developed. This model is grounded on (non-relativistic) QED and assumes that the operating principle of a microcolumn relies on the strong coupling of the ZPF to specific components that are found in neural tissue in very high concentrations. These components are molecules of the neurotransmitter type. We will prepare the ground for the theoretical description of the neurotransmitter-ZPF interaction and demonstrate that under the conditions observed in the brain it is plausible for a phase transition to occur. Such a phase transition gives rise to the formation of a coherence domain and entails coherence-triggered downstream effects that serve to regulate synaptic and axonal signal transduction, indicating that the neurotransmitter-ZPF coupling is crucial for the functioning of cortical microcolumns. This novel finding suggests that the large-scale activity patterns that are characteristic of high-level cognitive functions and make up the neural correlates of consciousness may be driven by a bottom-up process involving the ZPF. In this dynamic interplay, the microcolumns act at a mesoscopic level of organization that builds the bridge between the microscopic molecular level and the macroscopic level exhibiting long-range synchronization.

The structure of the article is arranged in such a way that Section II provides an overview of the basics of the brain and the current state of empirical evidence, both in terms of brain dynamics and brain architecture. In Section III, the physical model of a microcolumn is presented, followed by a discussion of the findings, culminating in the conclusions (Section IV). Finally, in Section V, an outlook is given on future directions of research.

## II. OVERVIEW OF THE BRAIN BASICS

### A. Bain dynamics and criticality

The exceptional performance of the brain is most evident in advanced cognitive processes, the distinctive marks of which are complex spatiotemporal patterns of activity in the cortex. It turns out that these patterns, which exhibit enormous variety and extend over large cortical areas, result from the reorganization of background activity [1] and reflect the cooperative behavior of a huge number of neurons [17]. A distinction must be made between perceptual processes, which are triggered by external stimuli and aim at experiencing the external world, and self-referential mental processes, such as stimulus-independent thought and memory retrieval. Perception occurs in rapidly forming frames with repetition rates that lie in the theta frequency band (8-12 Hz), with each frame corresponding to an *abrupt change in cortical dynamics* that leads to the formation of an attractor, the dynamics of which is characterized by synchronized activity in the beta (12-30 Hz) or gamma (30-50 Hz and beyond) frequency band [2, 3]. In self-referential cognitive processes, the attractor formations follow the alpha rhythm (4-8 Hz) [2]. There is growing neurophysiological evidence that the rapid reconfigurations of macroscopic brain dynamics resulting in the formation of attractors are due to *phase transitions* and that the organizing principle behind brain activity is based on criticality [2, 3, 5, 6, 17]. In this context, the concept of *self-organized criticality* is particularly important, which refers to the ability of a complex system to adjust a control parameter that allows the system to reside near a critical point and evolve toward a second-order phase transition [7].

A closer look at the processes underlying pattern formation reveals that the propagation of synchronized activity in cortical networks takes place in the form of *neuronal avalanches* whose sizes and lifetimes follow power law distributions, which are typical features of a system in a critical state and indicates that the dynamics of a critical system is dominated by universal properties [7, 18–20]. These avalanches are manifestations of the *collective organization* of spatiotemporal activity in the cortex [20] and display the periodicity of nested theta and beta/gamma oscillations [19]. A crucial finding is that the concentrations of the neurotransmitters glutamate and gamma-aminobutyric acid (GABA) as well as the main neuromodulators, such as dopamine, serotonin, and acetylcholine, have been identified as important control parameters for the regulation of neuronal avalanches [7, 21], with nested oscillations arising from excitatory-inhibitory networks that rely on the release of glutamate and GABA [22].

These insights point to the *pivotal role of neurotransmitters in inducing phase transitions*, an inference that is supported by studies which combine neurophysiological measures of brain activity in various frequency bands with the measurement of neurotransmitter concentrations. It turns out that there are significant changes in neurotransmitter concentrations during cognitive tasks [23], that brain activity in the theta frequency band is correlated with the glutamate concentration [24], that glutamate and GABA are involved in the brain-wide synchronization of activity, and that the glutamate concentration varies with the functional connectivity between brain areas [25]. Finally, also calculations based on phenomenological models designed to study phase transitions in cortical networks highlight the key function of neurotransmitters by showing that self-organized criticality is a phenomenon produced by synaptic dynamics [26], and by exposing that the emergence of avalanches is controlled by synaptic resources [27].

### B. Brain architecture

The dynamical properties of the brain are intimately connected with its architectural characteristics, particularly with the structural layout of the cortex (see Figure 1), which occupies a large part of the brain volume. The cortex is organized horizontally in layers and vertically, i.e., perpendicular to the cortical surface, in columns. The common view is that the basic unit of operation of the mature cortex is the minicolumn, also termed *microcolumn*, serving as a model of cortical organization [28, 29]. Even though there are differences between individual microcolumns in terms of structural details and connectivity, their basic design is the same throughout the cortical surface. A typical microcolumn comprises roughly 80 to 100 neurons, with little variation in size across species and its diameter ranging between 20 *µ*m and 60 *µ*m [29]. Notably, in the process of evolution cortical expansion has been achieved by steadily increasing the number of cortical microcolumns while retaining their size [28].

**FIG. 1.**
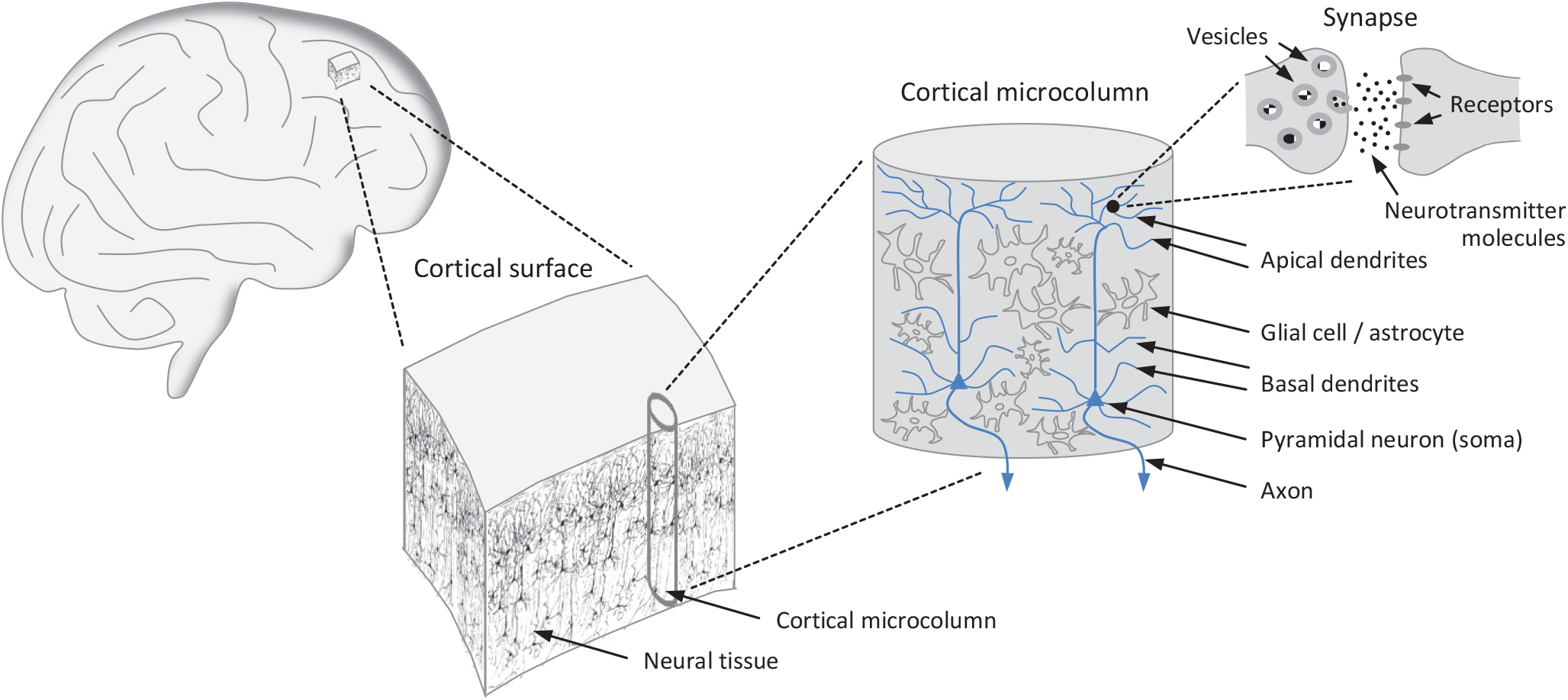
Organization of the cortex. The cortical surface consists of a vast number of microcolumns, which can be regarded as basic functional modules of the brain featuring uniform design principles. Essential components of the microcolumns are the pyramidal neurons, which are equipped with branched dendritic trees covered with synapses through which the neurons receive their inputs signals. The presynaptic terminal contains vesicles rich in neurotransmitter molecules, the release of which leads to the activation of receptors located on the postsynaptic terminal. The outputs of the neurons are transmitted via axons.

Substantial experimental support for the microcolumn hypothesis derives from the periodicity of the optical density of neural tissue [30]. Specifically, a repeating microcolumnar pattern consists in the bundling of apical dendrites of pyramidal neurons, which constitute roughly 80% of all neurons, with these bundles reaching an average diameter of approximately 30 *µ*m [31]. Moreover, the neurons of a microcolumn exhibit a significant degree of synchronized activity, suggesting that microcolumns constitute a brain-wide system of repeating functional modules [32, 33]. Finally, a high correlation is observed between dendritic and somatic activity, indicating that the pyramidal neurons within microcolumns function as integrated building blocks [34].

The microcolumns arrange themselves in larger associations, which in turn cluster into modality-specific areas, such as the somatosensory or the visual cortex. The columns are strongly interconnected with each other, and there are also connections to subcortical structures, in particular to the thalamus. Bundles of afferent fibers from cortical and thalamic modules project directly to the basal and apical dendrites of pyramidal neurons. These dendrites are densely covered with tens of thousands of excitatory, mostly glutamatergic, synapses through which inputs are received. The output channel of a pyramidal neuron is an axon, enabling it to connect to a large number of other neurons in adjacent or more distant microcolumns [28]. Another cell type found in microcolumns comprises interneurons, which are predominantly inhibitory and regulate the activity of pyramidal cells via GABAergic synapses [29]. The interplay of glutamatergic and GABAergic neurotransmission, along with the layered organization of the cortex, is crucial for the generation of oscillatory network activity [22].

Since in the following *we aim to achieve a basic understanding of the functional principle of an individual microcolumn*, we will disregard oscillatory network activity and ignore the layered architecture of the cortex as well as the presence of interneurons and GABAergic neurotransmission. This translates into a simplified structural model of a microcolumn as illustrated in Figure 2A.

**FIG. 2.**
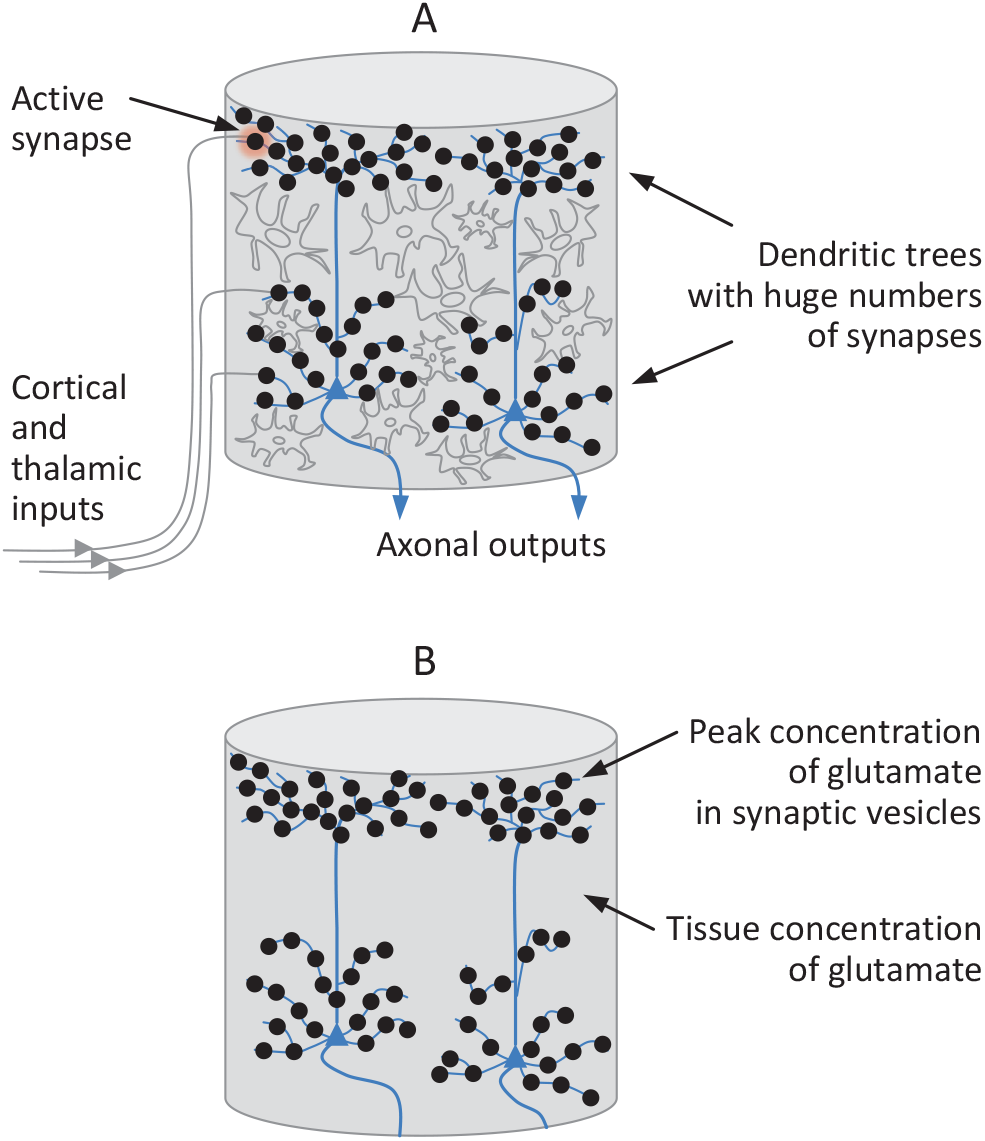
Structural model of a cortical microcolumn. (A) A microcolumn receives numerous inputs from other cortical areas and the thalamus. The cortico-cortical and thalamocortical fibers connect to synapses that populate the dendritic trees of the pyramidal neurons. In active synapses, signal transmission is effected by the release of neurotransmitters. A simplified model of a microcolumn includes only glutamate as an excitatory neurotransmitter. (B) In a further simplified model of an individual microcolumn the glial cells, fulfilling primarily a regulatory function, have been removed and replaced by effective glutamate concentrations, namely, a peak concentration localized in the synaptic vesicles and an average tissue concentration. According to this model, a microcolumn is viewed as a bundle of pyramidal neurons encased in a glutamate pool.

In view of the functional model to be developed in the following section, we take a closer look at excitatory neurotransmission. The most abundant excitatory neurotransmitter is glutamate, the concentration of which is several times higher in the brain than in other cell types and also higher than the concentration of any other molecular component in neural tissue, except water, with peak concentrations found in synaptic vesicles [35]. Glial cells regulate the glutamate pool and play an important part in the glutamate-glutamine cycle, which is a major metabolic flux in the cortex [35, 36]. More specifically, astrocytes, the most frequent type of glial cells, maintain glutamate homeostasis by controlling the balance between glutamate uptake and release, a process that is regulated by metabotropic glutamate receptors [37–39]. These regulatory processes are the basis for a tissue concentration of glutamate that is stabilized around a mean value.

This leads us to a further simplified model of an individual microcolumn, displayed in Figure 2B, which includes, other than pyramidal neurons, only glutamate as an excitatory neurotransmitter and relies on two glutamate concentrations, namely, a peak concentration localized in the synaptic vesicles and an average tissue concentration. In concrete terms, in this model the glial cells, fulfilling primarily a regulatory function, have been removed and replaced by effective glutamate concentrations. In this picture, *a microcolumn consists of a bundle of pyramidal neurons encased in a glutamate pool*. More precisely, the glutamate pool is to be understood as a *glutamate-water matrix* in which water plays an important supporting role.

## III. FUNCTIONAL MODEL OF A CORTICAL MICROCOLUMN

### A. Development of the model

Drawing on the foundations laid so far, we are now in a position to develop a functional model of an individual microcolumn. This model is predicated on the hypothesis that microcolumns are basic functional units of the brain that exploit the interaction of neurotransmitter molecules with the QED vacuum. Consequently, the model is based on a field-theoretical description of a many-body system interacting with the ZPF [40–42], which has previously been identified as an adequate approach for the treatment of brain dynamics [9]. More specifically, the power of such an approach lies in explaining the origin of abrupt phase transitions and understanding the collective behavior found in neural activity as a macroscopic feature of quantum origin [9–11].

The essence of the model is to describe the dynamical evolution of the many-body system under study, which in our case is a large ensemble of neurotransmitter (glutamate) molecules, not by looking at the dynamics of each individual molecule, but by focusing on the occupation numbers of the different molecular states. This allows us to achieve a holistic description of the ensemble of molecules using a *macroscopic wave function*. In the following, the corresponding QED-based formalism will be developed, leading us to equations expressing the dynamical evolution of the coupled neurotransmitter-ZPF system. The development of the formalism is guided largely by the seminal work of Preparata [41], as well as by works in which the approach has been applied to studying the properties of water [43, 44]. SI units are used in all calculations.

#### 1. Neurotransmitter-

The vector potential 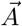 of the free electromagnetic field is given by a sum of normal modes

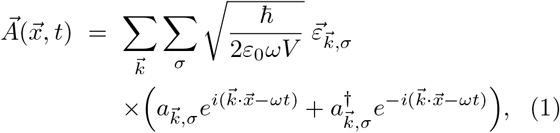

where *ħ* = *h*/2*π* is Planck’s constant, *ε*_0_ the vacuum permittivity, *V* the normalization volume, 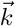 the wave vector, *ω* = 2*πν* the frequency, *σ* = 1, 2 the polarization index, 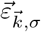 are the polarization vectors, and 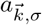 the field amplitudes, one for each dynamically independent degree of freedom of the field. The polarization and wave vectors satisfy

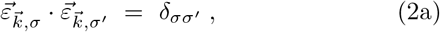

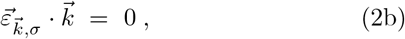

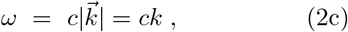

with *c* denoting the speed of light. The field amplitudes, which in the following are taken to be time-dependent, span a Hilbert space and obey the commutation relations

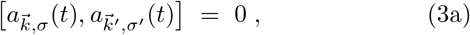

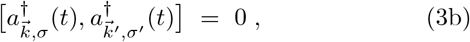

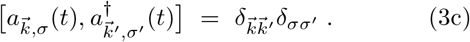

The eigenvalues of the number operator 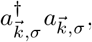 are to be understood as the occupation numbers of the corresponding field modes, fully characterizing a given field configuration.

In the Coulomb gauge 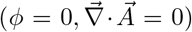, the electric field 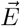 can be written as

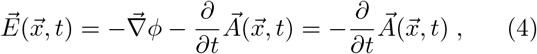

while the magnetic field 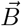 is given by

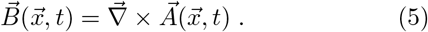

Using these equations as well as the commutation relations from Eqs. (3), the Lagrangian of the free electro-magnetic field can be expressed in the following form:

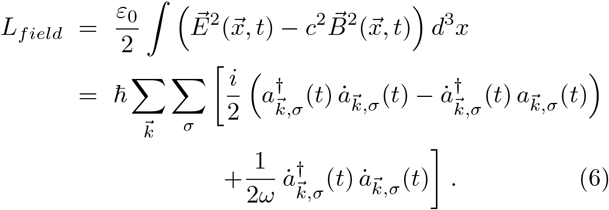

For the description of the matter system, which is taken to be bosonic and assumed to consist of N identical molecules, we use the wave function 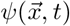, obeying the normalization 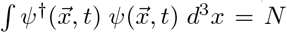. Only the glutamate molecules incorporated in the water matrix are taken into account here, as they dominate the interaction with the ZPF. All the other matter components of a microcolumn, such as molecules of the plasma membrane or receptor proteins of the pyramidal neurons, have too low concentrations to be significant for the resonant coupling to the ZPF (the function of these components will be discussed in Section IV). Later, we will write *ψ* as a linear superposition of molecular eigenfunctions, so that any configuration of the matter system, in analogy to the radiation field, is expressed by the numbers of quanta that populate the molecular eigenstates. Regarding the relevant eigenstates of glutamate, we can restrict ourselves to the *vibrational* states, since electronic excited states cannot be reached for energetic reasons and rotational states are frozen in the glutamate-water matrix, meaning that *ψ* represents a complete set of vibrational states in the electronic ground state of glutamate.

Employing the single-molecule Hamiltonian *H*_0_ and the short-range interaction Hamiltonian *H*_*SR*_, the full Lagrangian representing the matter system, the electro-magnetic field (ZPF), and the matter-ZPF interaction can be written as

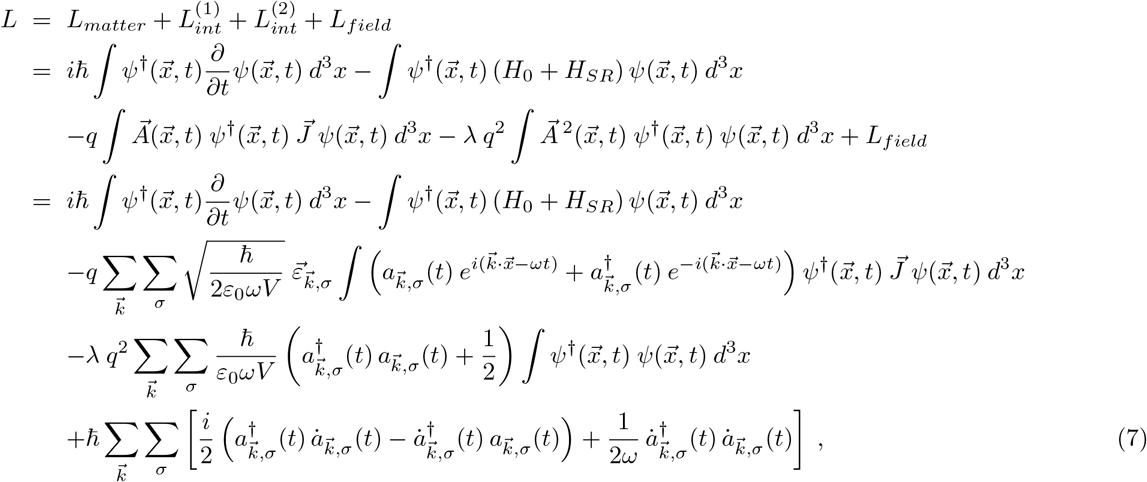

where *q* is the effective charge of the molecules (due to their polarization and dipole moment) and 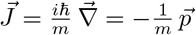, occurring in the first-order interaction term, stands for the current operator, with 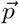 being the momentum operator and *m* being the effective inertial mass of the oscillating molecules. The factor λ in the second-order interaction term will be discussed in more detail later.

As one can show [41], the dynamical evolution of a system in the large N limit is determined by the classical Euler-Lagrange equations. From the Lagrangian given by Eq. (7), we can therefore straightforwardly derive the evolution equations that describe the macroscopic wave function of the system 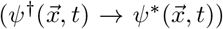 and the field amplitudes 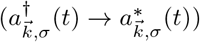 as follows

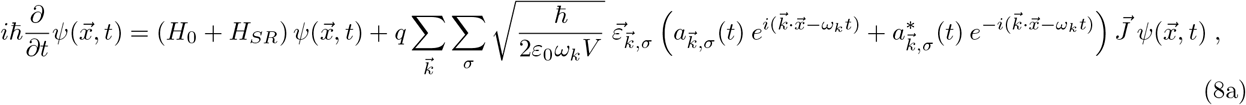

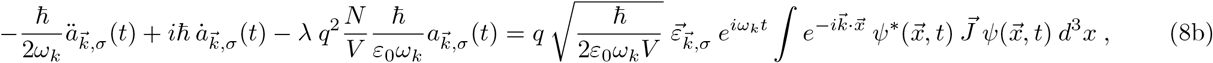

where we have explicitly indicated the *k*-dependence of the frequency (*ω*_*k*_) and exploited the normalization of *ψ*.

Even though the dynamics of the system is represented by the classical equations of motion, it is important to emphasize that Eqs. (8) are predicated on a *fully consistent quantum-theoretical treatment* of the coupled matter-ZPF system. This is evident from the fact that the ZPF, i.e., the presence of vacuum fluctuations, is a concept of quantum field theory that has no classical counterpart. Furthermore, in contrast to a semiclassical approximation, the amplitudes of the matter field and the radiation field obey commutation relations. The derivation of Eqs. (8) exploits the insight that in many-body quantum systems corrections to the classical path are suppressed by a factor 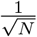, so that in the large *N* limit the dynamical evolution of the system is dominated by the classical equations of motion, which under suitable conditions lead to a stationary solution. Quantum corrections can be calculated systematically through perturbation theory [41], resulting in small corrections to the stationary solution while preserving its basic characteristics. In what follows, we are interested precisely in the basic characteristics of the stationary solution and the conditions under which it can form.

Decomposing the wave function *ψ* into a complete set of molecular (vibrational) eigenfunctions *φ*_*n*_ with energy eigenvalues *E*_*n*_

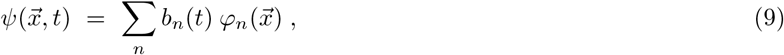

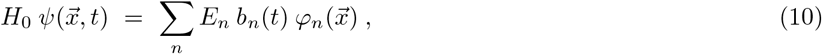

and rescaling the field amplitudes

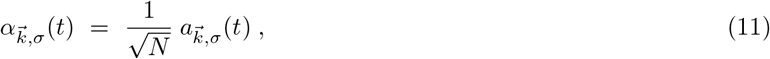

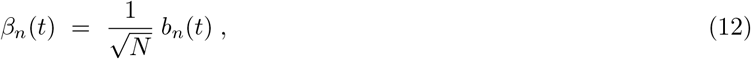

such that 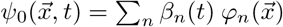 is normalized to 1, the evolution Eqs. (8) can be written as

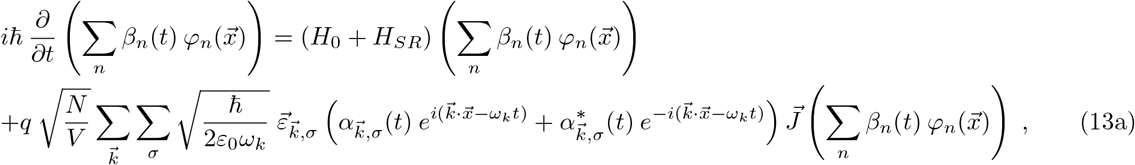

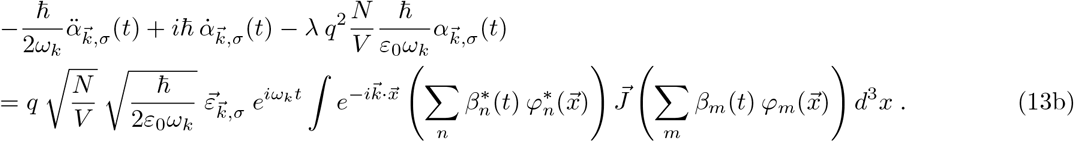

As one can see, the coupling constant q is amplified by a factor 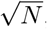, which already indicates the collective character of the matter-ZPF interaction. Therefore, even if the initial configuration of the radiation field corresponds to the perturbative ground state, it is to be expected that the strong matter-ZPF coupling will cause the perturbative ground state to be unstable and result, on the one hand, in the enhancement of particular field amplitudes and, on the other hand, in the excitation of collective oscillations of the matter system.

To further simplify the equations, we anticipate a result of the calculations performed below. These calculations reveal that the dynamical evolution of the coupled matter-ZPF system depends on highly selective resonance conditions which cause one of the molecular excited states to be singled out, subsequently termed *preferred excited state*, and the evolution of the system to be dominated by those ZPF modes that resonate with this preferred state, subsequently referred to as *dominant field modes*. Accordingly, the selective coupling of the radiation field to the ensemble of molecules gives rise to a situation in which only the amplitudes of those field modes are boosted that are in resonance with the preferred two-level transition. All the other excited energy levels not involved in the resonant matter-ZPF interaction can be disregarded without affecting the solution of the equations of motion (if we were to keep these energy levels here, they would drop out in later calculation steps). It should be noted, however, that the factor λ, as we will see in Eq. (30), includes the complete set of energy levels. Therefore, it is absolutely justified to restrict the sum over the energy levels in Eqs. (13) to the ground state with energy *E*_0_ and the preferred excited state with energy *E*_1_, where *E*_1_ − *E*_0_ = *ħω*_0_. Neglecting for the time being *H*_*SR*_ and switching to the interaction representation

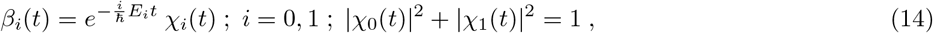

the evolution Eqs. (13) take the following form:

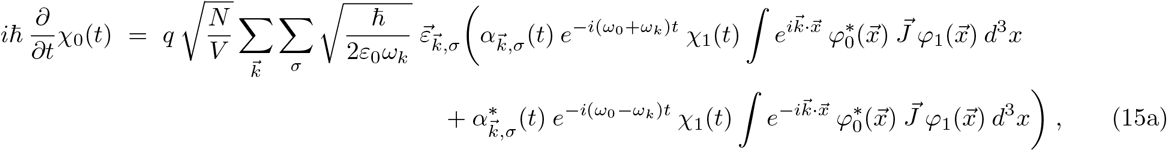

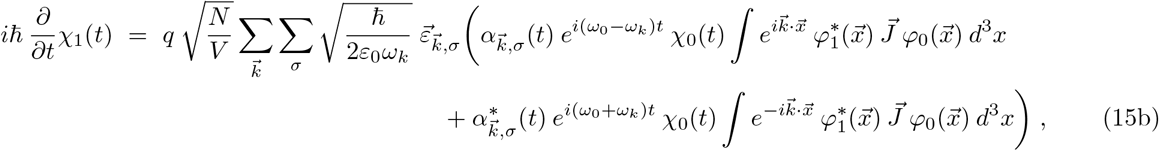

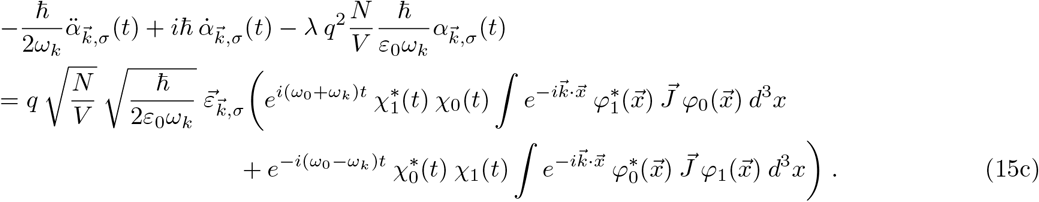

Since we are interested in the evolution over longer periods of time, we can apply the rotating wave approximation, which consists in neglecting time-oscillating factors. Taking the resonance condition *ω*_*k*_ = *ω*_*0*_ into account, yields

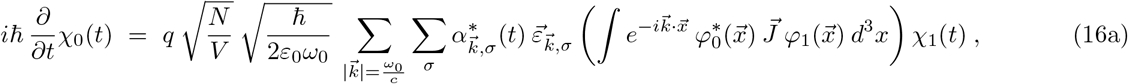

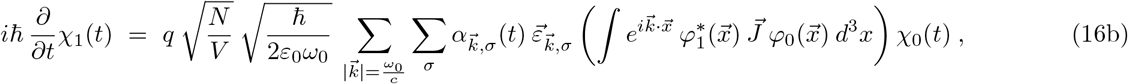

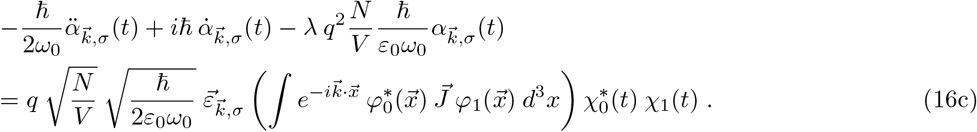

In the next step, we move to the dipole approximation, 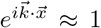, which is justified because the wavelength of the field modes involved in the interaction is significantly greater than the dimension of the individual molecules. Furthermore, we choose a compact notation for the transition matrix elements,

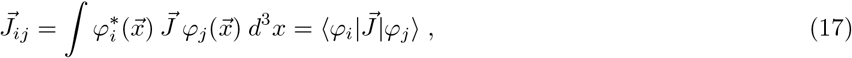

and write the sum over the wave vectors as an integral over all spherical angles, 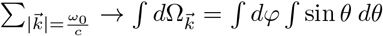 sin *θ dθ*, so that Eqs. (16) can be expressed as

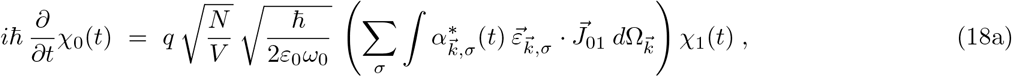

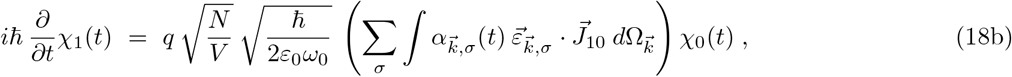

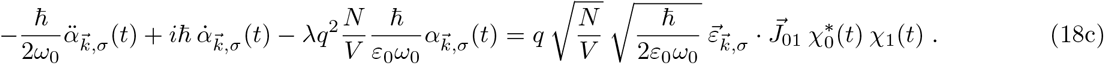

Setting 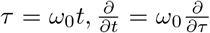, and in the following understanding all time derivatives as derivatives with respect to *τ*, yields

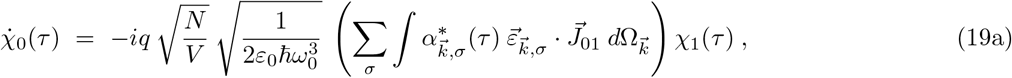

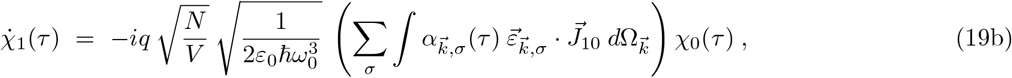

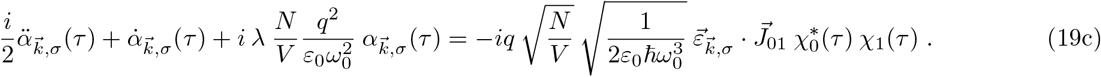

Introducing the integrated electromagnetic field amplitude of the dominant ZPF modes,

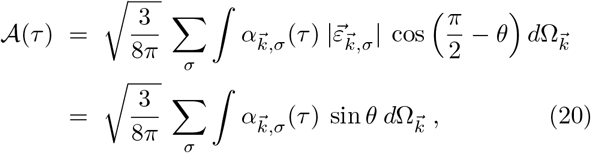

and defining two dimensionless quantities

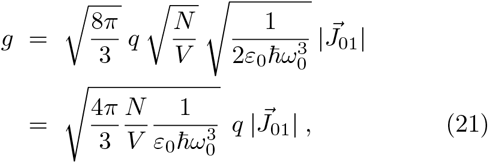

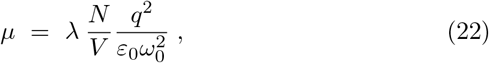

the evolution Eqs. (19), using the symmetry 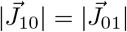, can finally be rewritten as

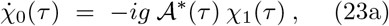

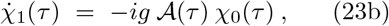

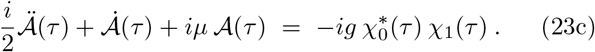

Thus, the dynamics of the molecules and the ZPF is described by a system of three coupled differential equations in which *χ*_0_(*τ*) and *χ*_1_(*τ*) represent the occupation numbers of the molecular ground state and preferred excited state, respectively, and 𝒜(*τ*) denotes the effective amplitude of the electromagnetic field modes that are in resonant interaction with the molecules. The parameter *g* plays the role of a coupling strength.

Instead of using the differential equations to calculate the time evolution of the system, the main focus of the following analysis will be on the two essential phases of the evolution, namely, the initial stage and the stationary state. The exact evolution between the two phases is not decisive for the derivation of the conclusions.

#### 2. Dynamics in the initial (runaway) stage

Using Eqs. (23) and taking into account the initial conditions *χ*_0_(0) ≈ 1 and *χ*_1_(0) ≈ 0, the early phase of the dynamical evolution, which we will also refer to as the *runaway stage*, has to obey the condition

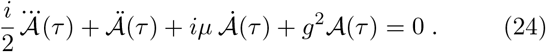

Looking for solutions of the form 𝒜(*τ*) = *C e*^*ipτ*^, we get the equation

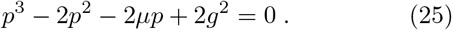

An exponential growth of the field 𝒜 occurs if this third-order polynomial has one real and two complex conjugate solutions, which is the case if 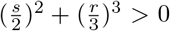, with 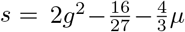 and 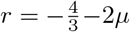. This leads to the following *runaway criterion*

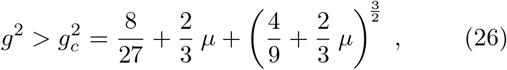

with *g*_*c*_ denoting the *critical coupling strength*.

In preparation for later calculations, we take a closer look at the quantities *g* and *µ* that go into Eq. (26). Expressing the current operator 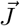 by the Hamilton operator *H*_0_ and the position operator 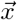,

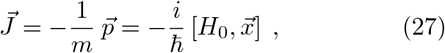

Using 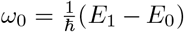, and switching over to the dipole operator 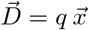, we get

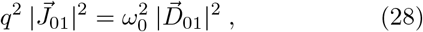

so that Eq. (21) can be rewritten as

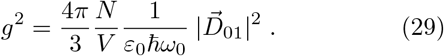

In order to calculate *µ*, we must address the quantity λ. A complete treatment of the interaction of the matter system with the radiation field using second-order perturbation theory, which requires the inclusion of all second-order terms in the interaction Hamiltonian [41, 45], yields

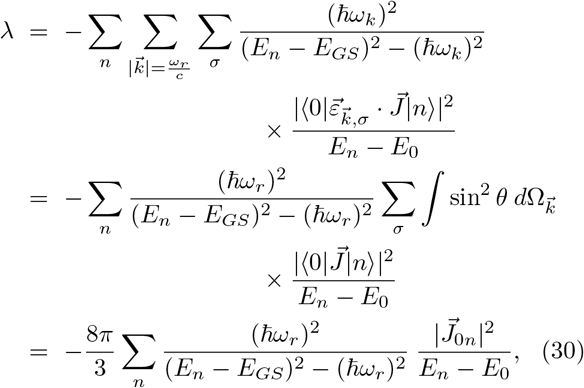

where *ω*_*r*_ denotes the resonance frequency, *E*_*GS*_ is the energy of the ground state, and *n* runs over a complete set of vibrational eigenstates |*n*⟩ with energy *E*_*n*_. Setting *ω*_*r*_ = *ω*_0_, *E*_*GS*_ = *E*_0_, 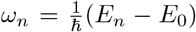, and using 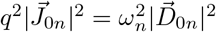 according to Eq. (28), we can rewrite Eq. (22) as

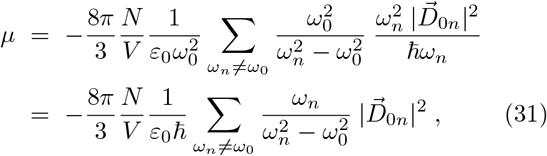

where the excited state with the resonance frequency *ω*_*0*_ (carrying the index *n* = 1) has to be excluded from the sum [43].

#### 3. Stationary solution of the evolution equations

To examine the stationary state of the coupled matter-ZPF system, we return to Eqs. (23). Considering the normalization condition for the occupation numbers, see Eq. (14), a suitable ansatz for solving the differential equations is given by

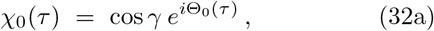

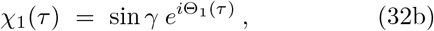

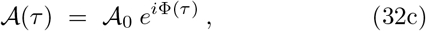

with 𝒜_0_ being real and positive. It can be shown that the solutions must take this form in order to satisfy the required conservation laws, in particular probability, energy and momentum conservation, demonstrating that the ansatz provided above describes the stationary state we are looking for [41]. The achievement of a stationary state gives rise to the formation of a *coherence domain* [46, 47].

Inserting Eq. (32) into Eqs. (23) reveals that a consistent solution has to respect the constraint

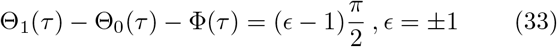

and requires 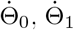 as well as 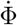 to be independent of *τ*, with

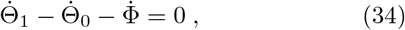

resulting in the following expressions:

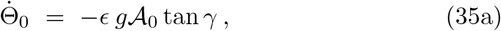

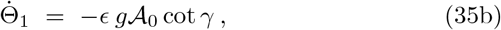

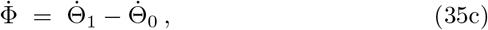

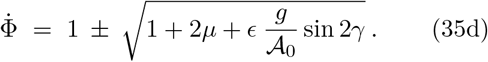

Two conserved quantities of the stationary solution are the dimensionless momentum 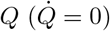 and the dimen-sionless energy 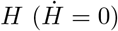 [41], which are given by

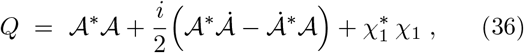

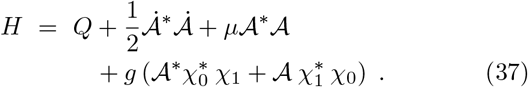

Employing Eq. (32) and setting *Q* = 0, one obtains

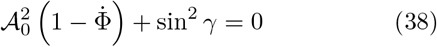

and

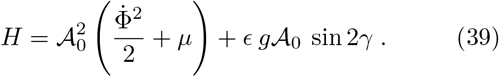

From Eq. (38) we see that 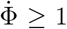 must hold, so that in Eq. (35d) the solution space with the plus sign has to be selected. On the other hand, it follows from Eq. (39) that in order to get the solution with the minimum energy, the condition *ϵ* = − 1 must be satisfied. Thus, the stationary state, which depends on the five parameters 𝒜_0_, γ, Θ_0_, Θ_1_, and Ф, is completely determined by the following set of equations:

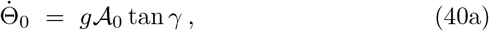

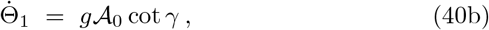

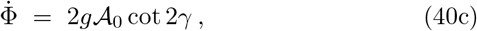

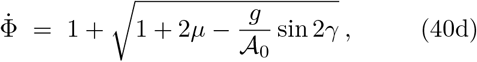

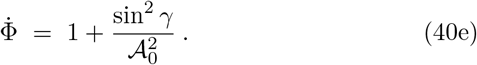

Eq. (39) can be further transformed by utilizing

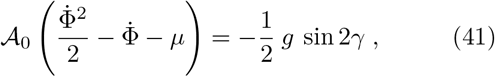

which results from Eq. (23c), and by exploiting Eqs. (40d) and (40e), leading to

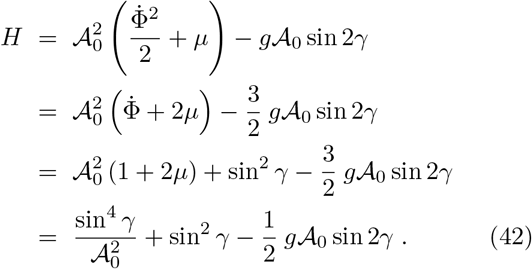

Looking at Eq. (42) in more detail, it turns out that under appropriate conditions (see later calculations in Sec. III B 2) the third term exceeds the sum of the first two terms, making *H* negative. Accordingly, compared to the energy per molecule in the perturbative ground state (*E*_0_), the energy per molecule in the coherent state (*E*_*coh*_) is reduced, with

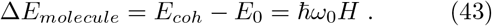

The formation of a coherence domain, as described by Eq. (40), thus corresponds to an *energetically favored state of the system* that is shielded by an *energy gap*

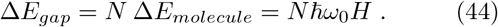

From the dynamical behavior of 𝒜, which is determined by the quantity Φ, it can be deduced that in the stationary state the electromagnetic field modes that are in resonant interaction with the molecules, being part of the spectrum of 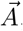, oscillate with the frequency

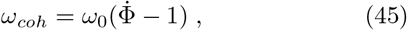

meaning that within the coherence domain the oscillatory dynamics is shifted from *ω*_0_ to *ω*_*coh*_.

In order to calculate *µ*, which enters Eq. (40d), we make recourse to Eq. (30), where we have to replace |0⟩ by |*coh*⟩ = cos *γ*|0⟩ + sin *γ*|1⟩, yielding

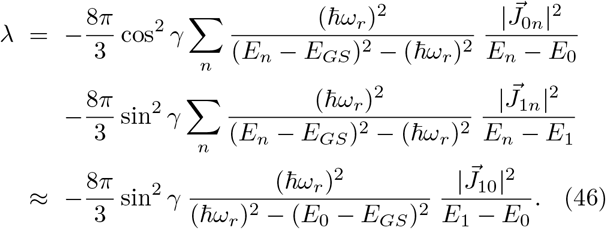

The approximation results from the sum over the states being dominated by one single term, which is a consequence of the shifted oscillatory dynamics in the coherence domain. Inserting Eq. (46) into Eq. (22), setting *ω*_*r*_ = *ω*_*coh*_, *E*_*GS*_ = *E*_*coh*_, using 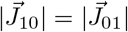, and exploiting Eqs. (28), (29), (43) as well as (45), we get

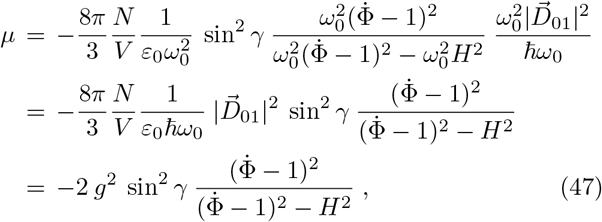

from which, using Eq. (40d), we can derive the condition

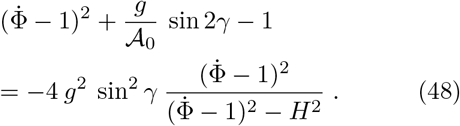

Thus, we can fully describe the dynamics of a coherence domain by calculating the coupling strength g according to Eq. (29) and finding a self-consistent solution for the parameters 𝒜_0_, *γ*, 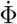, and H that satisfies Eqs. (40c), (40e), (42), and (48). The extent of a coherence domain is determined by the wavelength of the dominant field modes. The exact calculation can be found in Appendix A, with the result that, using *ω*_0_ = 2*πν*_0_, the diameter of a coherence domain is given by the relation

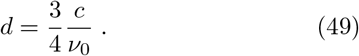

#### 4. Summary of the model

Starting from the structural model of a cortical microcolumn, which attributes an important part to the glutamate pool, we have derived evolution equations describing the interaction of a large ensemble of glutamate molecules with the ZPF. The evolution equations reveal that the amplification of selected ZPF modes occurs spontaneously when a specific runaway criterion is met. This criterion is tied to the magnitude of the glutamate-ZPF interaction, represented by a coupling strength that is determined by the concentration of the molecules and their excitability, which can be deduced from the dipole transition matrix elements. Upon exceeding a critical coupling strength, the strong interaction with the ZPF drives the ensemble of initially independent molecules toward a coherent state. The adjustable parameter of this process, which is also referred to as a superradiant phase transition [45, 48, 49], is the concentration of the molecules. Once the runaway stage has been triggered and a phase transition is in progress, the system undergoes reorganization and switches to a stable configuration in which the molecules and the selected ZPF modes oscillate coherently. This configuration is energetically favored and associated with a decrease in energy per molecule, resulting in the coherent state being shielded by an energy gap.

The selection of the ZPF modes involved in the reorganization of the system and the formation of the coherent state is based on resonance, which is determined by two factors, namely, first, the match of the frequencies of the field modes with the molecular excitation frequencies and, second, the strength of the dipole transitions between the molecular ground state and the excited states. These resonance conditions cause one of the molecular excited states (i.e., the preferred excited state) to be singled out and the evolution of the system to be dominated by those ZPF modes that resonate with the preferred excited state. In this way, the resonant interaction between the ensemble of molecules and the ZPF drives the entire system toward a stationary state that is characterized by the amplitude of the dominant field modes being significantly boosted and the molecules residing in a collective state. All the other field modes not involved in the resonant interaction remain in the perturbative ground state.

The achievement of a stationary state gives rise to the formation of a coherence domain.

Following these considerations, the postulated functional principle of a microcolumn can be divided into two steps. In the first step, illustrated in Figure 3A, the resonant glutamate-ZPF interaction triggers the *runaway stage* in individual clusters within the microcolumn, thereby inducing a local amplification of the dominant field modes. These clusters are the synaptic vesicles in which glutamate shows a *peak concentration*. Based on this first step, the initiation of a phase transition that pervades the entire microcolumn requires a second step, see Figure 3B, which consists in the simultaneous activation of many synapses, as only under this condition the individual runaway clusters will merge into a microcolumn-spanning cluster. More precisely, the release of highly concentrated glutamate from numerous synaptic vesicles distributed across the dendritic trees generates a single percolation cluster, which is the prerequisite for setting off an avalanche process that drives the glutamate pool within a microcolumn toward a stationary coherent state. The establishment of a stationary state entails the formation of a *coherence domain*, the dynamical properties of which are determined by the *tissue concentration* of glutamate and the diameter d of which is determined by the wavelength of the dominant ZPF modes.

**FIG. 3.**
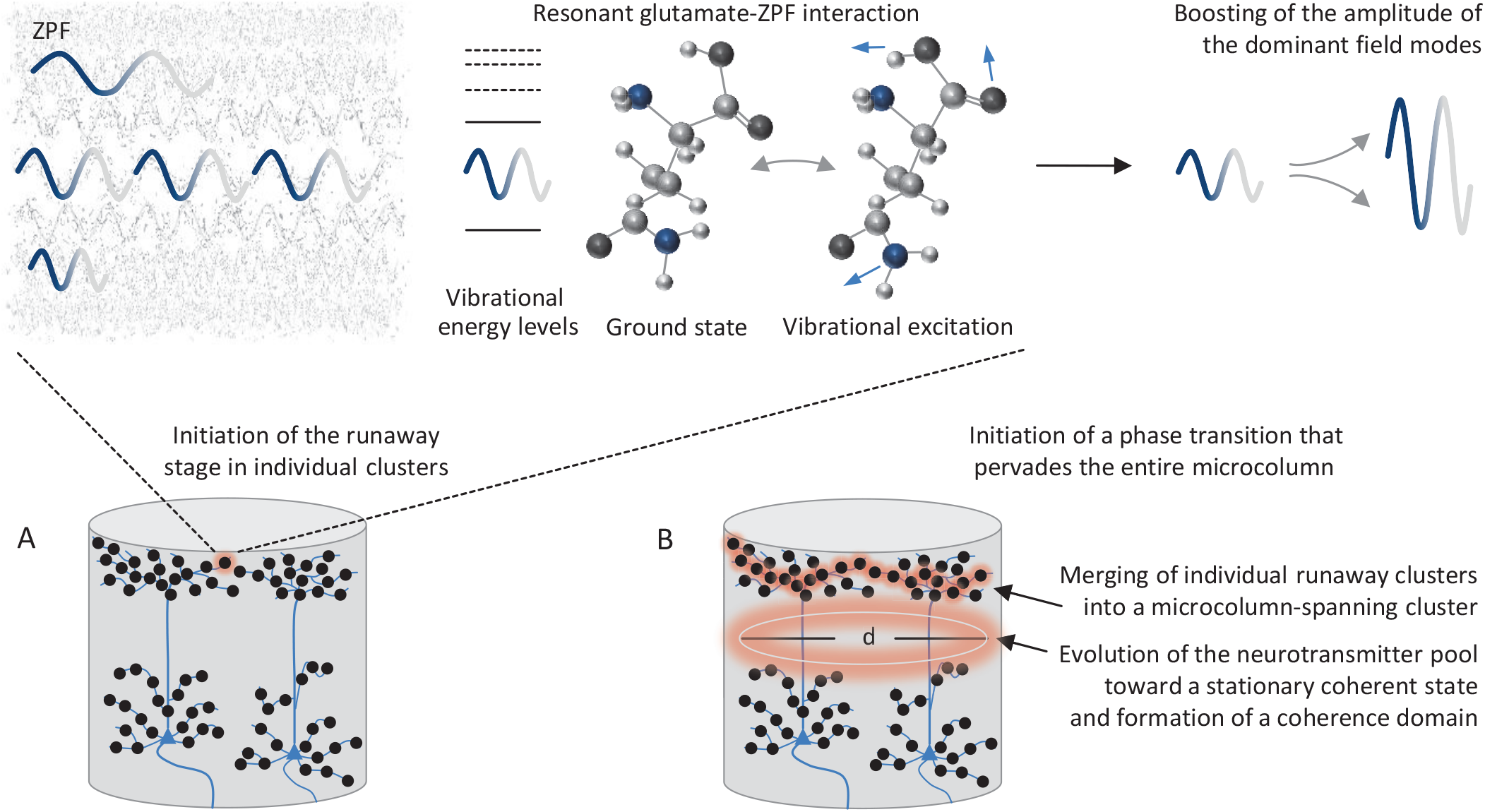
Postulated functional principle of a microcolumn. (A) The runaway stage in individual clusters (synaptic vesicles) is triggered by resonant glutamate-ZPF interaction. This interaction gives rise to the emergence of a collective state in which the amplitude of the dominant field modes is boosted and the molecules reside in a superposition of the ground state and the preferred excited vibrational state. (B) In order to initiate a phase transition that pervades the entire microcolumn, the simultaneous activation of many synapses distributed across the dendritic trees is required. Under this condition, the individual runaway clusters merge into a microcolumn-spanning cluster, setting off an avalanche process that drives the entire glutamate pool within a microcolumn toward a stationary coherent state. This process results in the formation of a coherence domain whose diameter *d* is determined by the wavelength of the dominant ZPF modes.

### B. Feasibility of the model

In the following, the feasibility of the postulated functional principle of a microcolumn and thus the plausibility of the presented model will be put to the test. In concrete terms, this means, on the one hand, that it must be studied whether the peak concentration of glutamate in synaptic vesicles is sufficient to initiate the runaway stage. On the other hand, it has to be demonstrated that, given the tissue concentration of glutamate, a stationary coherent state can be reached, which is the prerequisite for the establishment of a coherence domain. In this context, it also needs to be studied whether the diameter of a coherence domain is in agreement with the extent of a microcolumn, which is known from empirical data. In addition to these basic physical questions, the minimum number of activated synapses needed for the formation of a microcolumn-spanning percolation cluster would also have to be examined. However, this examination is beyond the scope of this article. In what follows, it is assumed that the microcolumn receives a sufficient number of synaptic inputs to form a percolation cluster.

#### 1. Initiation of the runaway stage

To start with, we turn to the runaway stage and look at the runaway criterion that contains the quantities *g* and *µ*, both of which depend on the dipole transition matrix Elements 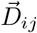. Since these matrix elements are difficult to calculate for neurotransmitter-type molecules, it is reasonable to obtain the corresponding values from empirical data, more specifically from the measured absorption spectra of the molecules. The relationship between the matrix elements and the experimentally measured absorption coefficients is derived in Appendix B. With the help of (B7), we can rewrite the expressions for g and *µ*, given by (29) and (31), arriving at

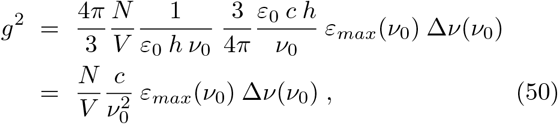

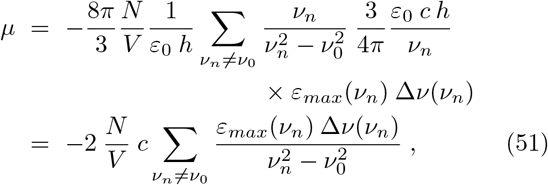

where *ε*_*max*_(*ν*_*i*_) denotes the peak values of the molar extinction coefficient (one value for each spectral line at res-onance frequency *ν*_*i*_), and Δ*ν*(*ν*_*i*_) stands for the widths of the spectral lines (full width at half maximum). According to Section III A 4, we have to plug in the vesicular glutamate concentration, while we extract the resonance frequencies, the widths of the spectral lines, and the peak values of the molar extinction coefficient from the available absorption spectra.

Regarding the neurotransmitter concentration in synaptic vesicles, values around 300 mmol/l are reported [50], with more recent findings on the vesicular glutamate concentration suggesting significantly higher values, reaching up to 700 mmol/l [51, 52]. For the further considerations, we will use the more conservative value and set

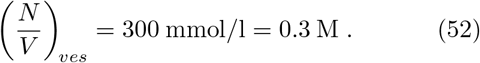

As with other biomolecules, the vibrational modes of neurotransmitters lie in the THz range and are due to the collective motion of intramolecular subunits as well as atomic groups that are involved in intermolecular interactions [53, 54]. For our calculations, it is crucial to note that the neurotransmitter molecules exist in an aqueous solution and are thus embedded in a kind of active water matrix, which can be attributed to water not being an inert solvent but, rather, forming hydration layers around solutes [55]. THz spectroscopy of water implies that this frequency band is dominated by highly correlated molecular motion, in particular by collective vibrations of hydrogen bonds, suggesting a significant influence on the dynamics of the solutes that engenders the amplification of vibrational resonances [55, 56]. More specifically, studies show that the enhancement of the absorption coefficients of the hydrated biomolecules depends on their hydrophilicity [56]. For glutamate, the most lipophobic neurotransmitter, which in aqueous solutions is ionized and found to form NaGlu ion pairs, it turns out that the effective dipole moment of the molecular anions embedded in the water matrix is considerably increased compared to the bare anions [57]. These findings are in good agreement with measurements according to which the molar extinction coefficients of hydrated NaGlu are substantially elevated relative to anhydrous glutamate [58].

In our field-theoretical approach, the amplifying effects induced by the water matrix are attributable to the previously neglected short-range interaction Hamiltonian *H*_*SR*_. Using in the following the absorption spectrum of hydrated NaGlu measured in the lower THz range [58] and taking the vesicular concentration from Eq. (52), we obtain

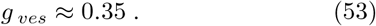

For the calculation of *µ*, the full absorption spectrum is required, for which in the case of glutamate, however, only limited information is available, since so far no data have been acquired for frequencies above 5 THz [58–60]. Therefore, in order to estimate *µ*, we resort to GABA, for which the broadband THz characteristics are known. In combination with the glutamate spectrum, the GABA characteristics provide at present the best possible insight into the absorption coefficients of neurotransmitters and the vibrational dipole transition matrix elements that derive from them, even though there is the limitation that the absorption spectrum of GABA has not been measured in an aqueous solution [61]. As the structure of the spectrum, in particular the spacing between the individual absorption lines, is a major factor in the determination of *µ*, this limitation is not expected to be crucial. From the analysis of the absorption spectrum, the lowest value for *µ*, namely

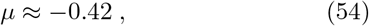

is obtained when the resonance frequency *ν*_0_ is equated with the absorption line at 7.8 THz. The resulting value pair for *g* and *µ* is displayed in Figure 4, including the curve that defines the critical coupling strength *g*_*c*_, see Eq. (26). As it turns out, *the value pair lies in the critical regime in which the runaway criterion for the initiation of a phase transition is fulfilled*.

**FIG. 4.**
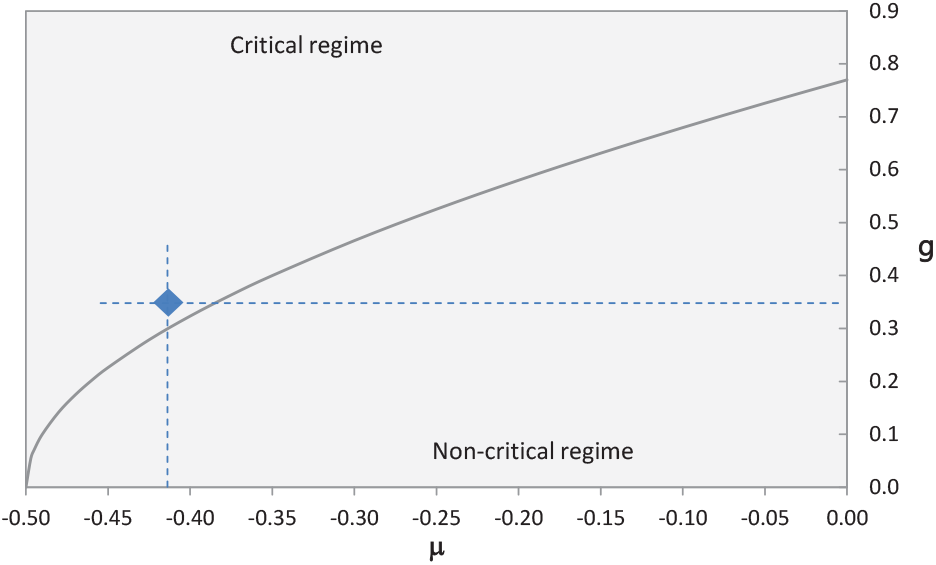
The solid curve indicates the critical coupling strength *g*_*c*_, see Eq. (26), as a function of the parameter *µ* and separates the critical from the non-critical regime. The blue diamond marks the pair of values (−0.42; 0.35) that results from the available data.

For the other absorption lines, the values of *µ* are below the critical limit, which gives the frequency of 7.8 THz a particular significance, in the sense that this is, on the one hand, the resonance frequency of the preferred excited state and, on the other hand, the frequency of the dominant ZPF modes that drive the evolution of the system. Throughout all of the subsequent considerations, we will therefore set

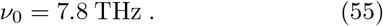

#### 2. Formation of a coherence domain

The essential prerequisite for the formation of a coherence domain is that, after triggering the runaway stage, the glutamate pool within a microcolumn is driven toward a stationary coherent state. More precisely, it has to be demonstrated that the evolution equations determining the dynamics of the coupled glutamate-ZPF system have a stationary solution. To this end, we return to the approach described at the end of Sec. III A 3, which consists in calculating the coupling strength g according to Eq. (29) and finding a self-consistent solution for the parameters 𝒜_0_, *γ*, 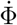, and *H* that satisfies Eqs. (40c), (40e), (42), and (48).

Following Sec. III A 4, we must now equate the neurotransmitter concentration with the tissue concentration. It has been known for many years that the glutamate concentration in brain tissue lies above 8 mmol/l [62]. More recent measurements of the glutamate concentration in different brain areas indicate values in the range between 8.4 and 14.6 mmol/l [24] or 7.7 and 17.1 mmol/l [63]. Based on these data, we choose a reasonable mean value of the glutamate concentration and set

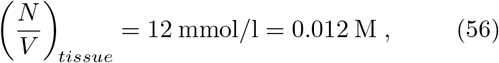

which is compatible with the average glutamate concentration in rodent cortex [64]. Using this mean concentration, the coupling strength *g* decreases to one fifth of the vesicular value, so that we get

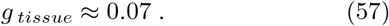

Applying numerical methods, *a stationary solution can be found*, which is specified by the following (approximate) values:

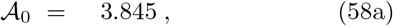

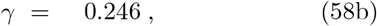

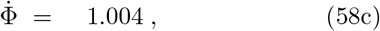

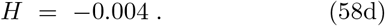

This solution can be interpreted to mean that, as a result of the strong glutamate-ZPF interaction, the amplitude of the dominant field modes is significantly elevated and the system is driven toward a collective state in which the glutamate molecules reside in a superposition of the ground state and the preferred excited vibrational state.

Using Eqs. (49) and (55), the diameter of a coherence domain can be estimated to be

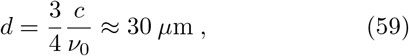

which is *well in accordance with empirically backed findings* on the extent of a microcolumn [29], in particular with the average diameter defined by the bundling of apical dendrites of pyramidal neurons [31].

Furthermore, it can be deduced from Eqs. (43), (55), and (58d) that in the coherent state the energy per molecule is reduced compared to the perturbative ground state, with

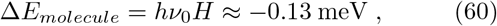

yielding a substantial energy decrease in the glutamate pool, Δ*E*_*gap*_ = *N* Δ*E*_*molecule*_ (see Eq. (44)), due to the immense number of molecules involved in the coherent state. Using the tissue concentration of glutamate, see Eq. (56), and the diameter of a coherence domain, see Eq. (59), the number of molecules involved amounts to *N* ≈ 10^11^. *The formation of a coherence domain thus corresponds to an energetically favored state that is shielded by a considerable energy gap*.

Finally, employing Eqs. (45), (55), and (58c), the shifted oscillation frequency within a coherence domain turns out to be

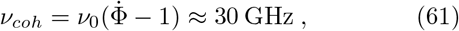

meaning that within a microcolumn the frequency of the dominant field modes, i.e., those ZPF modes that are strongly coupled to the glutamate molecules, is shifted from *ν*_0_ to *ν*_*coh*_ ≈ 30 GHz. This insight will play a role in the later discussion of the downstream effects resulting from the formation of a coherence domain.

#### 3. Considerations on decoherence

In the calculations made so far, disruptive thermal effects have been disregarded. The presence of such effects is commonly raised as an argument for the implausibility of the survival of coherent states in the warm and noisy environment of the brain [65]. However, a crucial finding of the presented model is that the formation of a coherence domain entails an energetically advantageous state that is isolated by a substantial energy gap. It is precisely the emergence of such an energy gap that, with a sufficiently large number of molecules involved, protects the collective state against thermal fluctuations [46].

The protection can be explained by the fact that within a coherence domain all the molecules of the glutamate-water matrix oscillate in unison with the dominant ZPF modes, which is why destructive thermal effects threatening the integrity of a coherence domain can only attack via its surface [42]. Therefore, the number *N*_*vul*_ of vulnerable molecules of a coherence domain that are subject to collisions with molecules from the environment and via which thermal energy can be fed into the system is significantly smaller than the total number *N* of molecules that constitute the coherence domain. Taking into account that the thermally vulnerable molecules are located only in the outermost molecular layers of a domain, we find that *N*_*vul*_ is of the order of 10^8^, which is a roughly a thousandth of *N*. Consequently, even though at a body temperature of 310 K the thermal energy per molecule is *E*_*th*_ = 26 meV and thus 200 times |Δ*E*_*molecule*_|, the inflow of thermal energy via the surface cannot easily overcome the energy gap, as *N*_*vul*_ *E*_*th*_ ≈ 0.2 *N* |Δ*E*_*molecule*_| < |Δ*E*_*gap*_|, resulting in markedly delayed decoherence. Moreover, the strong coupling of the molecules to the ZPF supports the self-preservation of a coherence domain.

Other research has also shown that *an energy gap is the key for the protection of coherence* in strongly interacting many-body systems [66], and that decoherence is highly suppressed in a large ensemble of particles forming a coherent state shielded by an energy gap [67]. A closer examination indicates that an exponential decay of coherence applies only to independent single-particle systems and many-body systems without interaction, while in strongly interacting many-body systems the nature of the decoherence induced by the coupling to a thermal bath is significantly altered [68, 69]. More precisely, the decay of coherence in systems with strong interactions follows a power law determined by a system- and interaction-specific decoherence time scale [68, 70]. According to model calculations, this time scale is proportional to the squared coupling strength as well as the squared particle number, implying that *decoherence is greatly slowed down in a system with strong coupling and a large number of particles* [70].

Furthermore, it should be recalled that the glutamate pool within a microcolumn is embedded in a water matrix. It has been found that liquid water, particularly interfacial water in close proximity to hydrophilic surfaces, is itself made up of coherence domains, so that the presence of water provides additional shielding from destructive thermal influences [71, 72].

All these considerations taken together *suggest that under the special conditions encountered in a cortical microcolumn*, i.e., the highly concentrated glutamate pool strongly coupled to the ZPF and embedded in a protective water matrix, *the formation and temporary maintenance of macroscopic quantum coherence is feasible*. Unequivocally proving that the survival time of the coherent state lies at least in the millisecond range and is thus sufficient to be neurophysiologically effective requires a calculation of the decoherence time scale, which will be addressed in a future work.

### IV. DISCUSSION AND CONCLUSIONS

In our model of a cortical microcolumn, we have so far been focusing on the pivotal role of the neurotransmitters, first and foremost glutamate. In concrete terms, the calculations have shown that the strong coupling of the glutamate molecules to the dominant modes of the ZPF drives the glutamate pool of a microcolumn toward a stationary coherent state. This supports the notion that the functional principle of microcolumns is based on macroscopic quantum coherence, explaining that microcolumns operate as integrated functional units of the cortex. In the following, it will be discussed that this functional principle, which is linked to the formation of a coherence domain, entails downstream effects that additionally substantiate the plausibility of the proposed model. These downstream effects can be divided into two categories, namely, effects caused by the coherent state of the glutamate molecules, and effects induced by the amplification of the dominant ZPF modes.

Regarding the first category, we have discovered that, as a result of the strong glutamate-ZPF interaction, the glutamate molecules are driven toward a stationary coherent state that turns out to be a superposition of the ground state and the preferred excited vibrational state. It is reasonable to assume that *the vibrational excitation of glutamate promotes conformational changes in glutamate receptors* located on postsynaptic terminals, leading to the opening of ion channels and *enhanced synaptic signal transduction*. This is in line with the principle of receptor activation through agonist-specific vibrational energy transfer that has already been examined in greater detail and identified as a promising approach to the understanding of agonist-receptor interaction [73, 74].

As far as the second category is concerned, our calculations have revealed that when a coherence domain is formed, the amplitude of the dominant field modes driving the evolution of the coupled glutamate-ZPF system is significantly elevated. Moreover, within the coherence domain the frequency of the dominant field modes is shifted to *ν*_*coh*_ ≈ 30 GHz, which lies in the microwave frequency range. In other words, the resonant glutamate-ZPF interaction within a microcolumn gives rise to a strong *intra-columnar microwave radiation field*. Since biological membranes are known to be very sensitive to electromagnetic fields in the microwave frequency range, the membrane-field interaction could induce a phase transition that influences the membrane permeability [75], which is consistent with the hypothesis that resonant interaction between microwaves and specific coupling sites in ion channels modulates ion flows across the membrane [76]. This hypothesis is supported by experimental evidence demonstrating that microwaves regulate voltage-gated ion channels [77] and stimulate collective membrane oscillations, which can be proven to be of non-thermal nature [78]. Further studies indicate that microwaves facilitate signal propagation along neurons by altering the membrane permeability and increasing transmembrane ion flows [79], and that microwave radiation has a direct effect on voltage-gated membrane channels of pyramidal neurons, thereby influencing the firing rate and the shape of action potentials [80]. These findings corroborate the notion that *the intra-columnar microwave radiation field plays the role of modulating voltage-gated ion channels and controlling axonal signal transduction*.

By exposing these mechanisms, the model is capable of explaining not only the high correlation between dendritic and somatic activity in pyramidal neurons [34], but also the observation that the pyramidal neurons of a microcolumn exhibit a significant degree of synchronized activity [32, 33]. The high level of synchronization within a microcolumn follows from the orchestrating role of the ZPF, which coordinates all those players inhabiting the sphere of influence of a coherence domain that are coupled to the dominant field modes. On the other hand, as no direct ZPF-based coupling is to be expected between microcolumns, synchronization among microcolumns can be achieved by means of ZPF-mediated controlled synaptic and axonal signaling.

Taken together, the body of evidence lends credence to the idea that the functioning of microcolumns is based on resonant glutamate-ZPF interaction and resultant macroscopic quantum coherence, which produces two types of downstream effects. These are the enhancement of synaptic signal transduction and the regulation of axonal signal transduction. The former effect is due to the vibrational activation of glutamate receptors located on postsynaptic terminals, while the latter effect derives from the modulation of voltage-gated ion channels populating the axonal membrane. Both effects combined may be crucial for the inter-microcolumnar communication and the formation of large-scale activity patterns that are characteristic of advanced cognitive functions.

From these considerations, the picture takes shape that *long-range synchronization in the brain emerges through a bottom-up orchestration process involving the ZPF, a key characteristic of this process being the formation, propagation, and synchronization of coherence domains*.

This insight opens up new vistas for our understanding of the fundamental mechanism underlying conscious processes [12–15], clearly distinct from alternative approaches that attempt to establish a link between quantum physics and consciousness [81–84].

## V. OUTLOOK

The presented model points to several research avenues that will be briefly outlined below.

The first path concerns the enhancement of the experimental data basis, specifically the provision of the broadband THz characteristics of hydrated NaGlu. The availability of these broadband characteristics would obviate the need for the fallback solution pursued here (see Section III B 1), which relies on the combined data from NaGlu and GABA, putting the calculation of the coupling strength of vesicular glutamate to the ZPF on an even more solid footing.

The model calculations have demonstrated that the stationary coherent state of the glutamate pool is isolated by a substantial energy gap, making it likely that this state is sufficiently protected against thermal noise to be neurophysiologically effective (see Section III B 3). It will be the subject of future studies to perform an estimation of the decoherence time scale.

Further research directions arise with regard to the downstream effects accompanying the formation of a coherence domain (see Section IV). On the one hand, this involves studying in greater detail whether vibrational excitations of neurotransmitters promote conformational changes in the matching receptors, initiating the opening of ion channels and enhancing synaptic signal transduction. On the other hand, a physical model should be developed to examine whether the modulation of ion flows across the membrane of pyramidal neurons is due to the resonant interaction between microwaves and specific coupling sites in voltage-gated ion channels.

The scope of the model presented here consists in disclosing the functional principle of an individual microcolumn. The long-term goal is to extend the model to the study of inter-microcolumnar communication and cooperative behavior that leads to the formation of long-range activity patterns and the establishment of oscillatory activity in the various frequency bands. Based on the extended model, it may be studied how synchronized activity patterns are influenced by changes in the concentrations of glutamate and other neurotransmitters (see Section II A). Furthermore, predictions regarding the dynamical characteristics of large-scale activity patterns should be strived for. These predictions are to be compared with dynamical key indicators derived from measurements of cortical dynamics, with the aim of lending further support to the notion of the ZPF being instrumental in the formation, propagation, and synchronization of coherence domains. If this route proves to be viable, the presented approach could turn out to be the bedrock of a fundamental theory of cortical dynamics.

## ACKNOWLEDGMENTS

This work was funded by the Stiftung Bewusstsein Kunst Kultur.

## Appendix A Extent of a coherence domain

Looking at the spatial structure of a coherence domain, we can assume spherical symmetry [41, 43] and write the electromagnetic field in the form

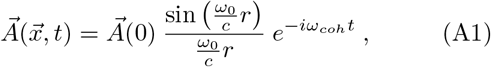

with *ω*_*coh*_ being determined by Eq. (45). In order to estimate the domain radius *r*_0_, we examine the radial components inside and outside of *r*_0_,

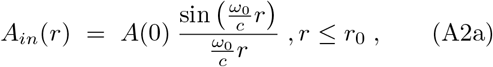

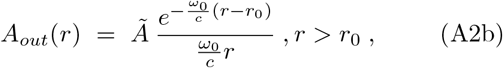

which must satisfy the two boundary conditions

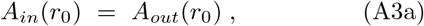

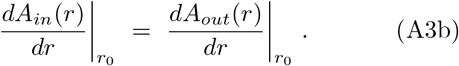

This translates into the constraints

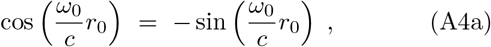

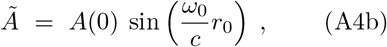

and results in the solutions

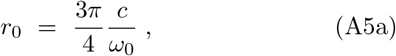

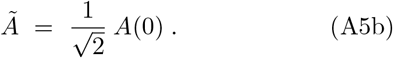

From Eq. (A5a), using *ω*_0_ = 2*πν*_0_, we obtain for the extent of a coherence domain the relation

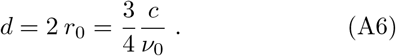

## Appendix B Determination of the transition matrix elements

The strength of the coupling between the pool of neurotransmitter molecules and the ZPF is determined by the dipole transition matrix elements 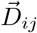, whose values can be obtained from empirical data, more specifically from the measured absorption spectra of the molecules. To this end, we derive a relationship between the matrix elements and the experimentally determined absorption coefficients [85]. In doing so, we concentrate on vibrational excitations of the molecules and look at levels *i, j* with energy *E*_*i,j*_ and occupation number *N*_*i,j*_, neglecting the degeneracy of the levels, since degeneracy arises only in the case of rotational transitions in the gas phase of the molecules. For the absorbed and emitted power related to transitions between the two levels we get

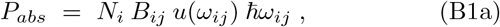

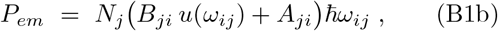

with *B*_*ij*_ denoting the absorption coefficient, *B*_*ji*_ the coefficient for stimulated emission, *A*_*ji*_ the coefficient for spontaneous emission, and *u*(*ω*) the spectral energy density. Assuming power balance in the thermal equilibrium, exploiting the symmetry *B*_*ji*_ = *B*_*ij*_, and equating the spectral energy density with the Planck distribution *u*(*ω*) = *ħω*^3^/[*π*^2^*c*^3^(*e*^*ħω*/*kT*^ − 1)], yields the relation

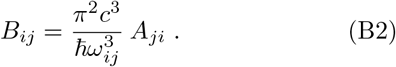

If we now consider the absorption of radiation in a medium of molecular concentration (*N*/*V*)_*med*_ = *n*_*med*_, the absorbed power per volume can be written as

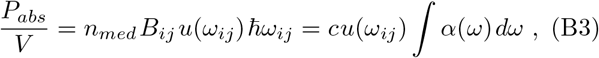

where we have introduced the frequency-dependent absorption coefficient *α*(*ω*) that carries the unit 1/m. For an absorption line at the resonance frequency *ω*_*ij*_, we thus arrive at

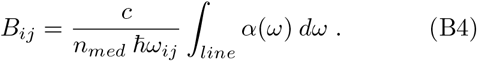

On the other hand, the power emitted by a molecule during a spontaneous dipole transition is given by [85]

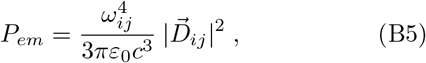

from which, using *P*_*em*_ = *A*_*ji*_ *ħω*_*ij*_ and Eq. (B2), we get

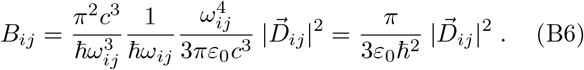

Utilizing Eq. (B4), we obtain for the dipole matrix elements the relation

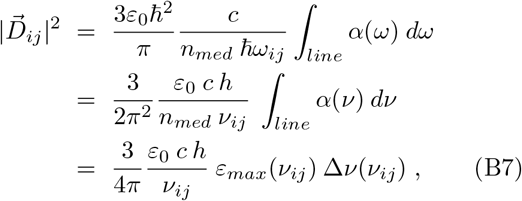

where we have introduced the molar extinction coefficient *ε*(*ν*) = *α*(*ν*)/*n*_*med*_ and exploited that the integral over a Lorentzian line shape can be represented as 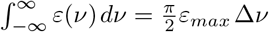, with *ε*_*max*_ being the maximum value of the corresponding absorption line and Δ*ν* being the full width at half maximum.

